# Genotyping by low-coverage whole-genome sequencing in intercross pedigrees from outbred founders: a cost efficient approach

**DOI:** 10.1101/421768

**Authors:** Yanjun Zan, Thibaut Payen, Mette Lillie, Christa F. Honaker, Paul B. Siegel, Örjan Carlborg

**Author notes:** Corresponding author: Örjan Carlborg, Department of Medical Biochemistry and Microbiology, Uppsala University, BMC, Box 582, SE-751 23 Uppsala, Sweden.

## Abstract

**Background:** Experimental intercrosses between outbred founder populations are powerful resources for mapping loci contributing to complex traits (Quantitative Trait Loci or QTL). Here, we present an approach and accompanying software for high-resolution genotype imputation in such populations using whole-genome high coverage sequence data on founder individuals (∼30×) and low coverage sequence data on intercross individuals (∼0.4×). The method is illustrated in a large F_2_ pedigree between lines of chickens that have been divergently selected for 40 generations for the same trait (body weight at 8 weeks of age).

**Results:** Described is how hundreds of individuals were whole-genome sequenced in a cost- and time-efficient manner using a *Tn5*-based library preparation protocol optimized for this application. In total, 7.6M markers segregated in this pedigree and 10.0 to 13.7% were informative for imputing the founder line genotypes within the F_0_-F_2_ families. The genotypes imputed from low coverage sequence data were consistent with the founder line genotypes estimated using SNP and microsatellite markers both at individual imputed sites (92%) and across the genome of individual chickens (93%). The resolution of the recombination breakpoints was high with 50% being resolved within <10kb.

**Conclusions:** A method for genotype imputation from low-coverage whole-genome sequencing in outbred intercrosses is described and evaluated. By applying it to an outbred chicken F_2_ cross it is illustrated that it provides high quality, high-resolution genotypes in a time and cost efficient manner.

## BACKGROUND

Genetic mapping of loci influencing the variation of quantitative traits (Quantitative Trait Loci, QTL) in intercrosses between divergent populations is a powerful strategy to dissect the genetic architecture of complex traits (Lynch and Walsh 1997). When produced from segregating founder populations, few of the polymorphisms at a particular locus in the intercross are fully informative about its founder line origin.

Therefore, statistical approaches have been developed to infer line origin from markers that segregate in the founder lines, e.g. (Haley *et al.* 1994; Crooks *et al.* 2011). The accuracy of the statistically inferred line-origin genotypes across the genome improves as the informativity and density of the genotyped markers increase (Haley *et al.* 1994). Nonetheless, genome-wide genotype inferences based on a few hundred markers limits the ability to capture individual recombination events. New genotyping strategies based on genome re-sequencing have been proposed and illustrated to deliver cost-efficient high-coverage genotyping to facilitate greater precision in estimation of the recombination events (Huang *et al.* 2009; Xie *et al.* 2010; Rowan *et al.* 2015; Campbell *et al.* 2018). These approaches were, however, developed for genotyping crosses between inbred lines. Further development to make them useful also in crosses between outbred lines would be of great benefit to, for example, researchers studying intercross populations from outbred lines of domestic animals and plants.

Here, we describe a cost- and time-efficient whole-genome, low-coverage sequencing approach to obtain genome-wide, high-density, and highly informative genotypes for studies of intercrosses between outbred founder lines. The properties of our method were illustrated by re-genotyping an F_2_ intercross (Jacobsson *et al.* 2005; Wahlberg *et al.* 2009) between the divergently selected body-weight Virginia lines (Dunnington and Siegel 1996; Márquez *et al.* 2010; Dunnington *et al.* 2013). Results demonstrate that this method delivered high-density and high-quality genome-wide genotypes where crossover events were identified with greater resolution and at a lower cost than that of reduced representation approaches based on a few hundred selected and individually genotyped genetic markers.

## RESULTS

### A new method and software for genotype imputation from whole-genome low-coverage sequence data

We developed a method (Figure 1) and software (https://github.com/CarlborgGenomics/Stripes) for genotyping crosses between outbred lines using whole-genome, low-coverage sequencing data.

**Figure 1.**
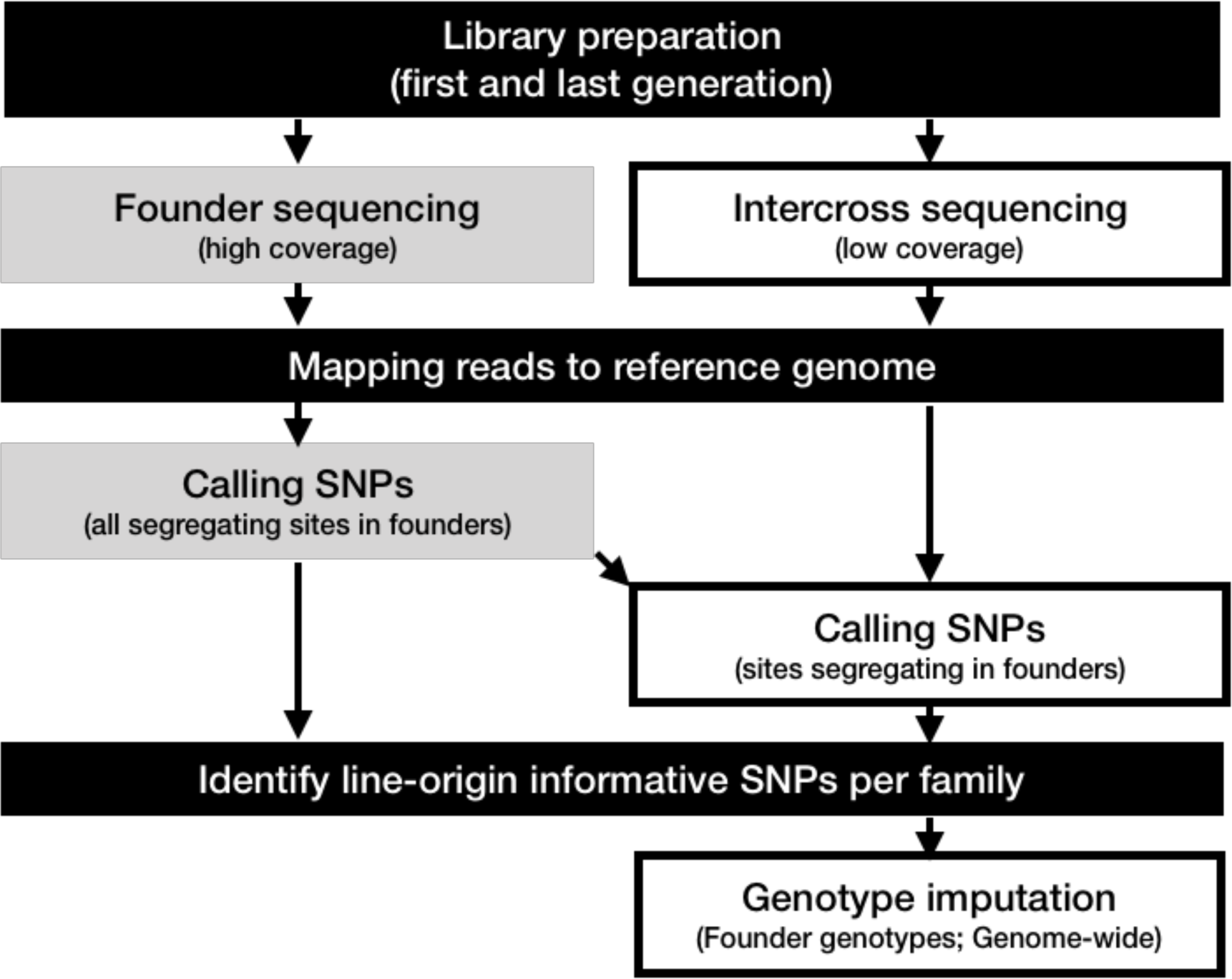
Outline of strategy for genome-wide imputation of founder-line genotypes in intercrosses between outbred founder populations. Black/grey/white boxes indicate analyses on the pedigree/founders/intercross individuals, respectively. Individual libraries for whole-genome resequencing are prepared for all individuals in the pedigree. For the intercross individuals, a high-throughput optimized version of a cost-efficient Tn5-based library preparation protocol originally developed by (Picelli, Björklund, Reinius, Sagasser, Winberg, and Sandberg 2014) is used. The founders from the divergent lines are sequenced to deep coverage and intercross individuals to low coverage. All SNPs segregating in the pedigree are identified using the founder sequences. Sequence reads on the intercross individuals are used to infer the chromosomal segments inherited from the founders using a modified version of a method originally developed for imputing genotypes from sequence data obtained on inbred crosses (Rowan et al. 2015).

The properties of the proposed genotyping approach were illustrated by analyzing data generated for an F_2_ population bred from the Virginia high (HWS) and low (LWS) body weight lines.

### Genome coverage by within full-sib family informative markers

In total, 7,608,483 SNPs were detected among the founders contributing to the evaluated F_2_ individuals (n_HWS_ = 27 and n_LWS_ = 29) using the high coverage (∼30×) individual sequence data. There were 213,946 markers fixed between the two founder lines and these were unevenly distributed across the genome (Figure S2). The numbers of markers fixed for alternative alleles between the HWS and LWS founders of the individual F_0_-F_2_ full-sib families (Within Family Markers, WFMrk) were considerably larger, on average 840,160 SNPs per family, leading to an average marker density of 791 WFMrk/Mb. The coverage of the genomes was relatively even, and on average 92% of the genomes were covered by more than 10 WFMrk/Mb in the evaluated families (Figure 2; Figure S3).

**Figure 2.**
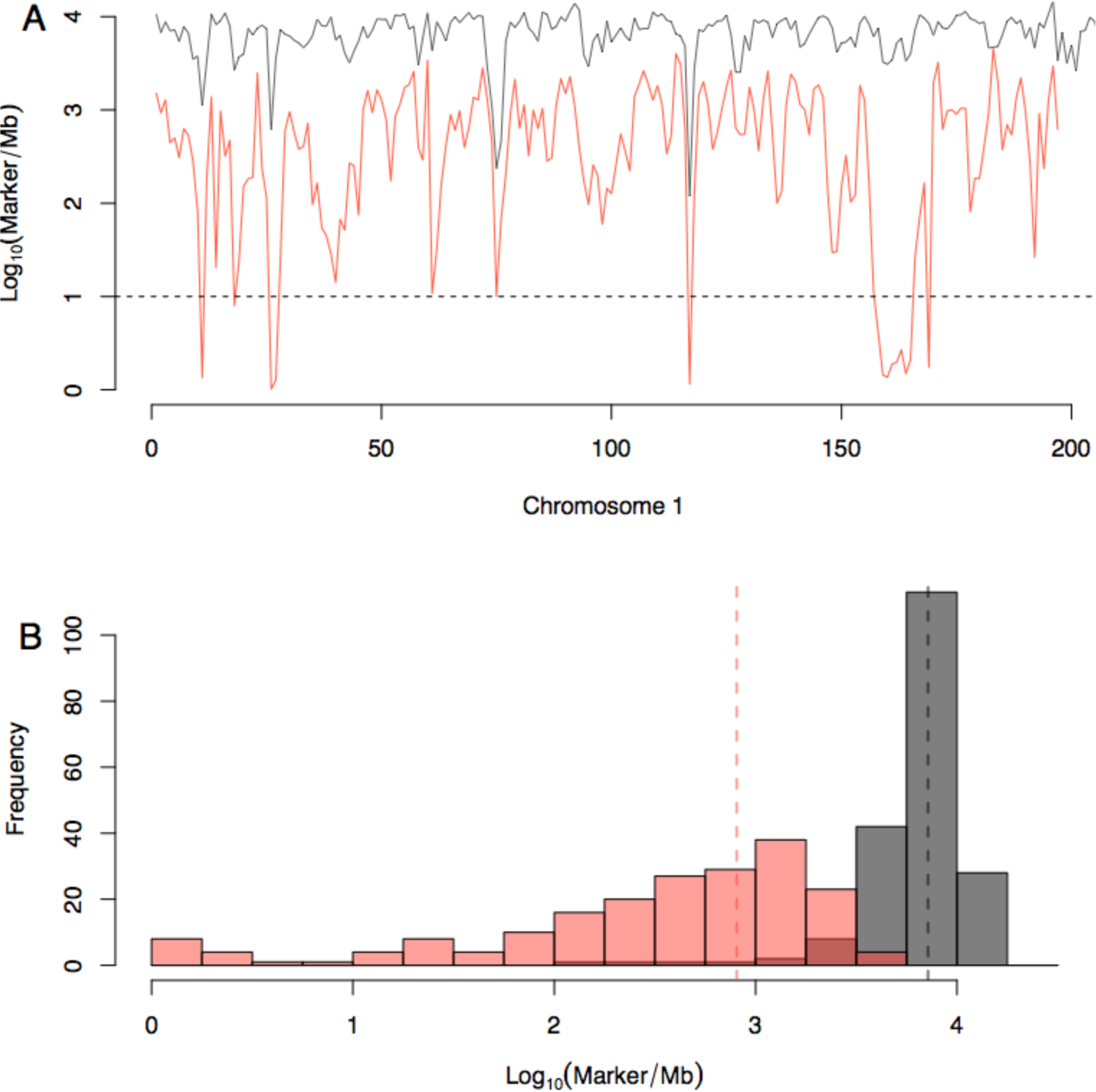
Illustrations of the density of markers fixed for alternative alleles between the HWS and LWS founders within the F_0_-F_2_families (Within Family Markers, WFMrk) on chromosome 1. **A)** Average number of markers in non-overlapping 1Mb bin along chromosome 1 (y-axis; log_10_transformed). The black/red lines represent the total number of markers segregating in the pedigree/the average number of informative markers in the 73 families in the Virginia lines F_2_pedigree. **B)** Distribution of the average numbers of WFMrk/Mb in the73 families (x-axis; log_10_transformed).

### Low coverage sequencing of intercross offspring

In total, sequence libraries were prepared for 837 F_2_ from the Virginia chicken lines intercross pedigree (Jacobsson *et al.* 2005; Wahlberg *et al.* 2009). 34 individuals failed in library preparation and another 14 individuals were removed due to low SNP marker coverage (<5 SNPs/Mb). The average sequence coverage for the 789 remaining individuals was 0.33′ resulting in an average coverage of 22.4 % of the WFMrk (188,520 on average) within the F_0_-F_2_ families.

### Genotype imputation from low-coverage sequence data in F_2_ offspring from outbred founder population

Genotypes were imputed for the remaining 789 F_2_ individuals with *TIGER* (Rowan *et al.* 2015) using data from sequence reads covering the WFMrk for each family. The raw imputed genotypes were filtered first by removing double crossover events within 3Mb (Figure S4) and next individuals with a genome-wide call rate less than 0.9 after crossover filtering. The final dataset remaining after this quality control included 728 F_2_ individuals.

### Quality of imputed genotypes

The quality of the imputed genotypes was assessed using the 728 individuals that passed our quality control and had genotype probabilities estimated from SNP and microsatellite data (Wahlberg *et al.* 2009). Overall, the genotypes for the evaluated non-overlapping 1 Mb bins were consistent for the two methods, both per bin in the population (on average 0.92 agreement; Figure 3A) and across the genome per individual (on average 0.93 agreement; Figure 3B).

**Figure 3.**
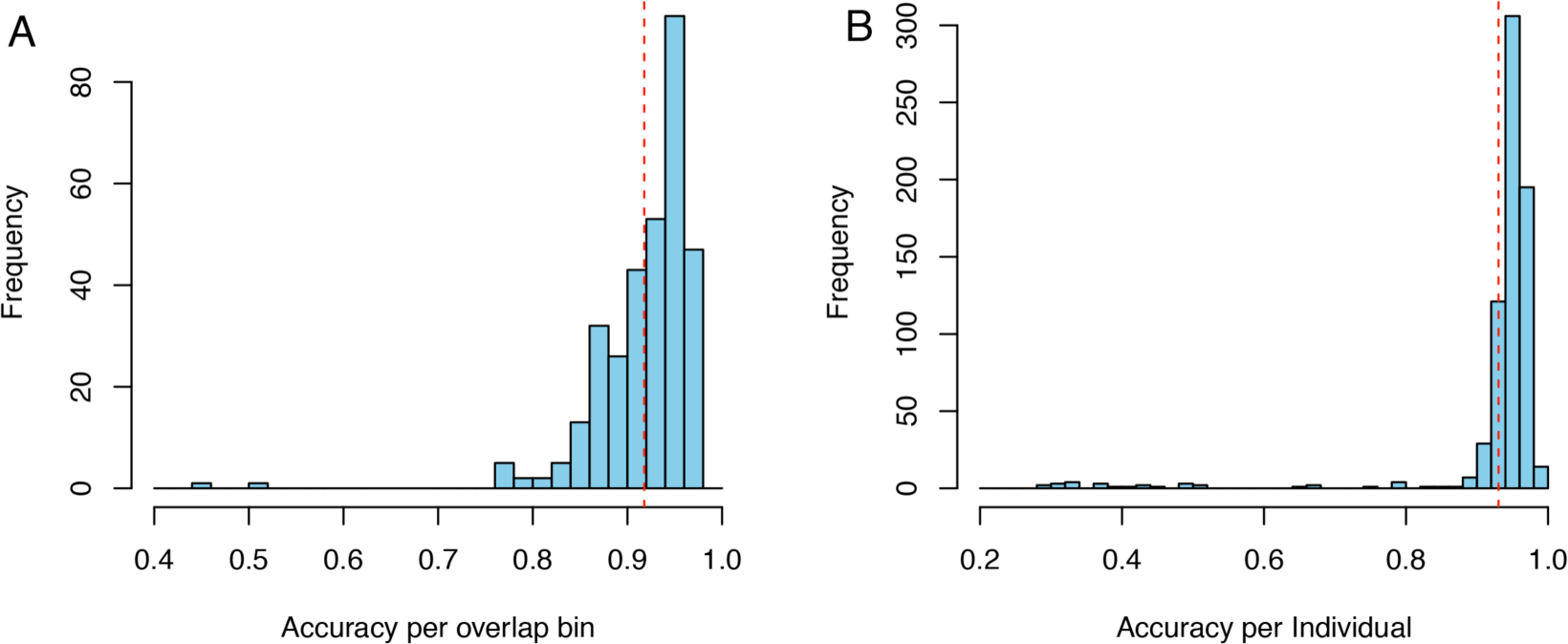
Estimated imputation accuracy in the Virginia lines F_2_population. Imputation accuracies were measured as the proportion of bins with the same genotype imputed from low-coverage sequence data and SNP/microsatellite data (Wahlberg et al., 2009) for **A)** the 322 1Mb bins across the genome and **B)** genome-wide across the 322 bins in the 728 evaluated individuals. Dashed red lines show the mean accuracies (0.92/0.93 for accuracy per bin/individual).

There was more variation in the accuracies per bin (Figure 3A) than per individual (Figure 3B). We evaluated if this could be because of differences in coverage among regions. First, the 14 individuals filtered out due to low average WFMrk coverage (<5 markers/Mb) were evaluated. For these, the sequence coverage ranged from 0.02-0.05×, and corresponding SNP densities were between 1.5-3.9 markers/Mb. Overall, the accuracies of the imputed genotypes were low in these individuals both per bin (0.38) and per individual (0.39). Next, the 789 F_2_ individuals passing our SNP coverage cutoff (> 5 markers/Mb) were evaluated. For these, the sequence coverage ranged from 0.03-0.86×, with corresponding SNP densities between 5.9-605.0 markers/Mb (Figure S3). There was, however, little effect of differences in coverage on imputation accuracy (Figure S3).

Figure 4 illustrates a comparison between the *TIGER* imputed genotypes and the genotype probabilities from Wahlberg *et al* (Wahlberg *et al.* 2009) for one F_2_ individual across chromosome 1. Overall, the chromosome wide genotypes were similar at 95.7% of the marker locations in Wahlberg *et al* (Wahlberg *et al.* 2009). In addition to this, the *TIGER* based imputation using the WFMrk provided imputed genotypes for many additional sites (Figure 4A vs 4B). As a result, the resolution of the recombination breakpoints increased considerably with 50% being resolved within <10kb (one example in Figure 4C). The higher marker density also allowed imputation of genotypes in areas that were not covered by the set of microsatellite markers selected by Wahlberg *et al* (Wahlberg *et al.* 2009). For example, this makes it possible to resolve heterozygous regions that are intermediate to homozygous regions agreed upon by both methods (blue in Figure 4B) and suggests longer putative double recombinant regions in the genome that were missed by the sparser marker set (yellow in Figure 4B).

**Figure 4.**
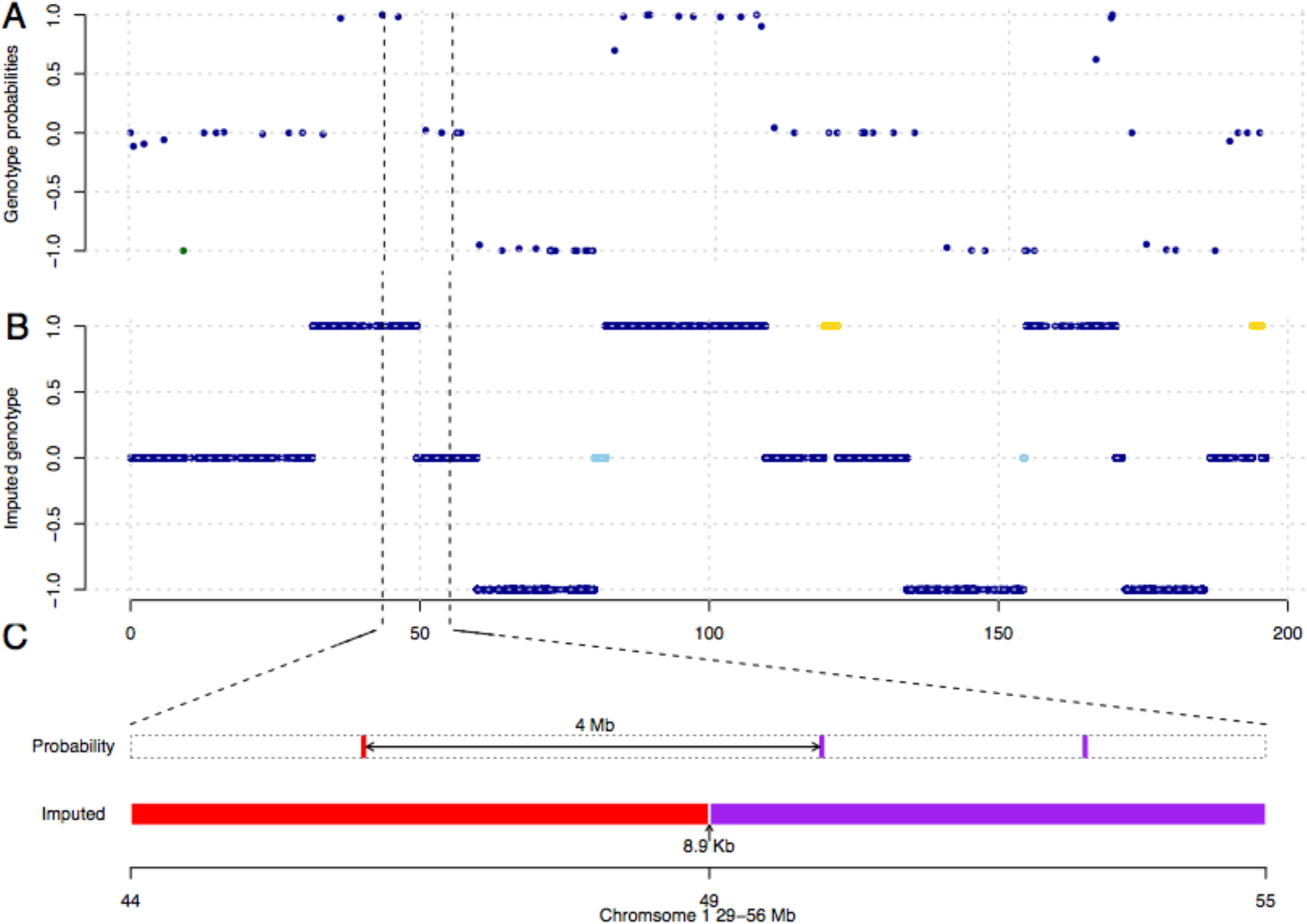
Resemblance of the genotypes imputed from low-coverage sequence data and genotype probabilities estimated from sparse individual genotypes across chromosome 1 for an individual F_2_ chicken. **A)** Genotype probabilities estimated by Wahlberg et al. (Wahlberg et al. 2009) using the algorithm described by Haley et al. (Haley et al., 1994) provided at the locations of the 74 SNP and microsatellite markers genotyped in this pedigree. The green dot illustrates a genotype that disagrees with those imputed from sequence data; **B)** Imputed genotypes from the low-coverage sequence data, where the color of the dots indicate overlapping genotypes by the two methods (dark blue), and expected heterozygous genotypes resolved (light blue)/novel putative double recombinant genotypes suggested (yellow) by imputing genotypes from sequence data; **C)** Illustration of the increased resolution from sequence based genotype imputation for one of the recombination breakpoints on chromosome 1 (between 44 – 55 Mb). Genotype probabilities inferred from SNP and microsatellite, and our pipeline are illustrated as colored bars, where red/purple indicate homozygous for the HWS allele/ heterozygous, respectively.

## DISCUSSION

Here, we describe a cost-efficient genotype imputation strategy from low-coverage whole-genome sequence data for experimental crosses between outbred founder lines. The founders of the pedigree were sequenced to a high coverage using standard approaches followed by a low cost whole genome sequencing protocol to facilitate low coverage sequencing of hundreds of intercross individuals (<10 €/individual including library preparation and sequencing). This is less than the cost of genotyping a few hundred individual SNP markers with existing strategies, even when accounting for the cost of high-coverage sequencing of the large (n = 56) number of founder birds in our pedigree. By utilizing pedigree information, genotypes can be efficiently imputed using strategies and statistical approaches previously developed for recombinant inbred populations (Huang *et al.* 2009, Rowan *et al.* 2015). In addition to decreasing the cost, there is a massive increase in marker density from a few hundred to several hundred thousand sites with imputed genotypes across the genome. In our dataset, 50% of the recombination breakpoints could be resolved to within 10 kb. Ultimately, this should enhance the development of linkage maps and increase resolution in linkage and QTL mapping. The resolution is, however, less than that achieved with inbred populations (Rowan *et al.* 2015) where individual recombination breakpoints in > 90% of the cases were resolved down to 2 kb resolution even at even lower sequence coverage (0.1×). This difference in resolution is most likely because of the smaller number of available founder line informative markers due to the genomic similarity of our intercross, which was founded from the same base population 40 generations prior to the intercrossing. Resolution could be increased by deeper sequencing, as only 22.4% of the markers were covered by reads at the current sequence depth, although several regions in the genome had a low number of informative markers, thus making it difficult to resolve even at higher sequencing coverage (Figure 2A).

For those individuals that passed quality control, there was good general agreement between the genotypes obtained via imputation from low coverage sequencing data and the genotype probabilities estimated from individual SNP and microsatellite markers. This agreement was of the same magnitude both per evaluated 1 Mb genome bin and across chromosomes 1-24 per individual. Although the genotypes were not identical at all evaluated sites, the average genotype imputation accuracies were 0.92 per site and 0.93 per individual. As illustrated in Figure 4, differences are likely to reflect situations where the higher coverage imputation strategy resolved intermediary genotypes in recombinant regions and likely double recombination events. The true genotyping accuracy of the imputation strategy is therefore likely to be significantly higher and sufficient for common analyses in F_2_ populations such as linkage and QTL mapping. It should be noted that genotype imputations are expected to be of lower quality in areas of the genome that are poorly assembled, as well as on smaller chromosomes/contigs/scaffolds where few informative markers exist.

In our data, no obvious increase in imputation accuracy was observed once the sequence coverage exceeded 0.05×. This suggests that sequencing to a lower depth than the currently applied 0.4× would have been sufficient to achieve acceptable imputation accuracy. The current maximum number of index combinations based on Illumina primer sequences that could be differentiated in 1 lane is 398. An attractive option for chicken (or other species of similar genome size) would therefore be to sequence the maximum number of individuals that could be multiplexed using these primers in a lane on the Illumina HiSeq4000 to obtain an average coverage of ∼0.20×.

The agreement between the genotypes scored by imputation from low-coverage sequence data and those estimated from non-fixed SNP/microsatellite genotypes was variable both per bin and per individual in our dataset (Figure 3). A possible explanation for this, suggested by the results shown in Figure 4, is that genotyping based on a few hundred selected markers will be less precise in regions where the genotype changes rapidly such as double recombinant regions or shifts from one homozygous to the other. A more detailed assessment of the contribution by this to the reported genotyping accuracies was, however, not possible from the available data.

The genetic divergence between the outbred founder populations used to breed the studied intercross needs to be considered when deciding how deep to sequence them in order to reveal sufficient number of markers for genotype imputations. Here, the founders were from two selection lines obtained by 40 generations of divergent selection from the same base population. Due to the lower expected genomic divergence between these lines (Figure 1; Figure S1) than in crosses between, for example, wild and domestic populations, we opted for a high (∼30×) coverage in this study to identify most of the SNPs segregating in the populations. We found that 0.03% of all identified SNPs were completely fixed for alternative alleles between the two founder lines and that 10.0-13.7% were fixed between the two pairs of F_0_ individuals used to breed each F_0_-F_2_ family. In populations where there is greater divergence between the founder populations, more markers will be informative. Maintaining this high sequence coverage of the founders is recommended as it will benefit both imputation accuracy as well as resolution of the recombination breakpoints by revealing more of the available informative markers.

In our quality control procedure, few (14; 2%) individuals were removed due to low SNP coverage, likely due to poor library quality or uneven pooling of libraries before sequencing. More problematic was the need to remove a larger number of individuals (n = 61 or 7%) due to F_2_ genotypes that were inconsistent with the genotypes of the F_0_ founders in the family. This might be the result of DNA/Sample mixup or contamination during sample processing. Ultimately, the individuals that passed quality control have high quality imputed genotypes illustrating the need and the value of the implemented quality control procedure to identify erroneous individual samples for further explorations.

In pedigrees where the founder populations are highly divergent, an alternative strategy for genotyping can be considered if it is desirable to minimize genotyping costs. Rather than sequencing individual founders to high coverage, we suggest pooling the libraries from the individual founders of each population and sequence the two pools to high coverage (∼30×) to identify markers that are fixed (or nearly so) between the populations. In our F_2_ population, employing this strategy would have reduced the number of SNPs useful for genotype imputation from 840,160 to 213,946. This number of genome-wide markers is sufficient for high-quality genotype imputation, but the uneven distribution of the divergently fixed markers in the genome would challenge the approach in some poorly covered regions (Figure S1). Our population is, however, likely represents an extreme case in this respect as the intercrossed lines are closely related and the number of founders is large. Despite this, the imputed genotypes based on these markers were of sufficient high quality in bins that are linked throughout the genome to facilitate efficient QTL mapping in the population (data not shown). When designing and genotyping an outbred intercross populations, many factors need to be considered. In this respect, it is worthwhile to note that while this pooled founder sequencing approach will be more efficient in populations where fewer and/or more distantly related founders are used, it is still likely to be sufficient for the purpose of building high-quality linkage maps and performing genome-wide QTL analyses even in populations with many founders from more closely related lines.

A common reduced representation genotyping strategy for intercross populations bred from divergent populations is to select a few hundred markers, that are either fixed for alternative alleles or segregating at highly divergent frequencies in the founders, for genotyping in the entire pedigree (Wahlberg *et al.* 2009). Selection of markers then requires prior information on which markers are polymorphic and informative for the differences between the populations. In contrast, our strategy is time effective, because no preselection of markers for genotyping is needed and facilitates a more complete exploration of the genome, as markers will be scored across all assembled regions and not only on the major chromosomes, scaffolds, or contigs.

### Conclusions

In summary, a method for genotype imputation from low-coverage genome sequencing in outbred intercrosses is described and evaluated. The results obtained from applying it to an outbred chicken F_2_ cross illustrate how it provides high quality, high-resolution genotypes in a time and cost efficient manner.

## METHODS

### Genotyping

Figure 1 outlines the developed genotyping by whole genome re-sequencing strategy for outbred intercross pedigrees. First, individual-based sequencing libraries (Zan and Carlborg 2018) were prepared followed by sequencing of the individuals from the two founder populations of the pedigree to high coverage for SNP calling (here 30×) and individual intercross individuals to low-coverage (here ∼0.4-0.8×). Genotypes for the intercross individuals were then inferred separately within each F_0_-F_2_ family: First, markers fixed for alternative alleles in the parents from the two founder populations were identified using the SNP calls from the high-coverage sequence data. Then, within-family offspring genotypes were inferred from low-coverage sequence data using the approach developed for inbred intercross populations (Rowan *et al.* 2015). All scripts used for SNP calling, genotype imputation, and downstream evaluations are publicly available on Github (https://github.com/CarlborgGenomics/Stripes and https://github.com/CarlborgGenomics/Stripes_downstream).

### Test dataset: A large F_2_ pedigree produced from the divergently selected body-weight Virginia lines

Data from a reciprocal F_2_ intercross between chickens from two divergently selected lines, obtained by bidirectional selection for body weight at 56 days of age (here, the high weight selected “HWS” and low weight selected “LWS”) (Dunnington and Siegel 1996; Márquez *et al.* 2010; Dunnington *et al.* 2013), were used to evaluate the proposed method. The base population for the lines was founded by crossing seven partially inbred lines of White Plymouth Rock chickens. The F_1_ population of the intercross was generated by matings of 10 males and 17 females from the HWS to 8 males and 22 females from the LWS. To generate the F_2_, 8 males and 73 females from the F_1_ were mated (Jacobsson *et al.* 2005; Wahlberg *et al.* 2009). Included in our analysis here are the 837 F_2_ individuals with complete F_0_-F_2_ pedigrees and DNA available for sequence library preparation, and the 56 founders (n_HWS_ = 27 and n_LWS_ = 29) that contributed to these F_2_ individuals.

### Whole genome sequencing of pedigree founders

Libraries for high coverage sequencing of pedigree founders were prepared using Illumina TrueSeq and sequences obtained by paired end sequencing (2 x 150 bp) on an Illumina HiSeq X (performed by the SciLifeLab SNP&SEQ Technology platform; Uppsala, Sweden). Mapping, SNP calling and quality control followed the Broad best practices. Reads were mapped to the chicken reference genome (galgal5; (Warren *et al.* 2017) using the Burrows-Wheeler Aligner (BWA–MEM v. 0.7.13 (Li 2014)).

Aligned reads were sorted and duplicate reads marked with picard (v. 2.0.1; https://broadinstitute.github.io/picard/). Base Quality Score Recalibration (GATK 3.7) was carried out, prior to SNP calling with HaplotypeCaller (GATK 3.7). Variants were filtered to MAF > 0.043, AC > 5 and QUAL > 30 resulting in a high quality set of SNPs were obtained for further analyses.

### Whole genome sequencing of intercross individuals

A *Tn5*-based protocol (Picelli, Björklund, Reinius, Sagasser, Winberg, and Sandberg 2014) for low-cost, high-throughput preparation of individual sequencing libraries (∼1€/library) was optimized for use in large-scale genotyping of the intercross individuals in the pedigree (Zan and Carlborg 2018). In short, genomic DNA was fragmentized and tagged using *Tn5* transpose purified from a plasmid available from AddGene (http://www.addgene.org/, pTXB1-Tn5; ID60240) (Picelli, Björklund, Reinius, Sagasser, Winberg, and Sandberg 2014). Dual indexes were attached during PCR amplification and subsequent size selection was performed using AMPure XP beads (Beckman: A63881). The detailed procedure for library preparation, pooling and quality control was described in (Zan and Carlborg 2018).

Sequencing of intercross individuals was performed using an Illumina HiSeq 4000. First, the two largest F_2_ full-sib families (n = 32) were sequenced to ∼0.8× coverage to test the quality of the prepared libraries and implemented pooling strategy (Oklahoma Medical Research Foundation Genomics Core). The remaining intercross individuals (n = 805) were then sequenced to ∼0.4× coverage by pooling ∼200 multiplexed individuals per lane (Texas A&M Genomics and Bioinformatics Service).

Demultiplexing of the dual indexed reads into individual fastq files and trimming of the adapters was done using *bcl2fastq* v2.17.1.14 (Illumina, Inc).

### Founder line genotype imputation in an outbred intercross pedigree

The low coverage sequence data from the F_2_ individuals were mapped to the *Gallus* V5.0 reference genome (Warren *et al.* 2017) using the Burrows-Wheeler Aligner (Li 2014). Only those reads where both pairs were uniquely mapped (mapping quality >=30) were retained. *Mpileup* in *BCFtools 1.8* (Li 2011) was used to extract the raw information at each polymorphic site (depth and SNP read). A custom Python script was used to reformat the output to contain only information about location of the polymorphism (chromosome and position), the genotype (reference and alternative allele call), and the read depth (https://github.com/CarlborgGenomics/Stripes).

To impute the genotypes across the genomes of the intercross individuals, we adapted an approach developed by Huang *et al* (Huang *et al.* 2009) for genotyping recombinant inbred lines derived from re-sequenced pairs of founders using low-coverage sequence data. First, the intercross population was divided into its component full-sib F_0_-F_2_ families where the F_2_ offspring share the same F_0_ founders. For each of these families, unique sets of SNPs were identified including those that were fixed for alternative alleles between the two F_0_ founders from HWS and the two F_0_ founders from LWS in each family. These sets were identified using a custom Python script (https://github.com/CarlborgGenomics/Stripes). The data could then be analyzed as if the population was inbred and line-origin genotypes could be imputed within families for each informative SNP using the *TIGER* software (Rowan *et al.* 2015). The imputed genotypes were compiled, using custom R scripts, into a recombination block (Rowan *et al.* 2015) genotype matrix for downstream analyses. In this matrix, each row represents one individual and each column a block in the genome in which no crossover event was identified.

The distribution of all the SNPs used to impute the F_2_ genotypes across the genome, as well as the distribution of the average number of SNPs that were informative in each F_2_ full-sib family, were summarized in relation to the genetic map used for a previous analyses of this chicken population (Wahlberg *et al.* 2009).

### Assessment of imputation accuracy

First, individuals with few called SNPs (genome wide average < 5 Markers/Mb) were removed from the dataset. This was because accurate genome-wide imputation requires a reasonably high marker density. Nearby double recombination events are biologically unlikely due to crossover interference and, in chickens, cytological evidence suggests it to be absolute in regions < 5 Mb on the largest chicken chromosomes (Zakharova *et al.* 2006). Raw *TIGER* (Rowan *et al.* 2015) imputed genotypes were filtered to remove double recombination events closer than 3 Mb as biologically these would be considered most unlikely.

Sample mix-ups, DNA contaminations, and pedigree errors in the data will lead to inaccurate genotype imputation. These errors will increase the number of inferred genome-wide crossover events in affected individuals. To filter out such individuals, we deleted samples where the genome-wide genotype call rate decreased to <90% after removing short (<3 Mb) double recombination events from the data. An alternative approach would be to remove individuals that were outliers in the distribution of genome-wide recombination events inferred in the genome using *TIGER* (Rowan *et al.* 2015), which would lead to a very similar final set of individuals (Figure S1).

The quality of the imputed line origin genotypes from the low-coverage whole-genome sequence data across chromosomes 1-24 were evaluated by comparing them to line origin genotype probabilities estimated at equidistant (1 cM) intervals across the genome in the same population from 434 individual SNP and microsatellite genotypes using the Haley and Knott algorithm (Haley *et al.* 1994) by Wahlberg *et al* (Wahlberg *et al.* 2009). In total, n = 728 of the individuals that passed our quality control filtering had genotype probabilities estimated in Wahlberg *et al* (Wahlberg *et al.* 2009) and could therefore be included in this evaluation. Comparisons were performed in 1 Mb bins across the evaluated chromosomes. For each bin, the imputed genotype - high weight homozygous (HH), heterozygous (HL), low weight homozygous (LL) – from the low-coverage sequence data were compared to the average genotype probabilities for the cM locations (Wahlberg *et al.* 2009) mapping to this bin. Bins with *TIGER* imputed recombination events were excluded from the comparisons. A genotype, HH/HL/LL, was assigned for a bin from the (Wahlberg *et al.* 2009) data if the average probability for a particular genotype exceeded 0.8. Otherwise, it was considered missing. The proportions of genotypes in agreement between the two methods per bin and across the chromosomes per individual were used as measures of genotyping accuracy (https://github.com/CarlborgGenomics/Stripes_downstream).

## DECLARATIONS

### Ethics approval

All procedures involving these animals were carried out in accordance with the Virginia Tech Animal Care Committee animal use protocols.

## Acknowledgements

Genome sequencing of the founders of the chicken pedigree was performed by the SNP&SEQ Technology Platform in Uppsala, which is part of Science for Life Laboratory at Uppsala University and is supported as a national infrastructure by the Swedish Research Council (VR-RFI). Protein production and purification was performed at the Protein Science Facility at Karolinska Institutet, Stockholm. Leif Andersson provided genotypes and samples for the F_2_ population. Special thanks are given to the multitude of students and colleagues involved in the development of these populations and phenotypic data obtained over the decades when the selected, relaxed, and intercross populations were reproduced and maintained at Virginia Tech in Blacksburg, VA. Without them and continuous support from Virginia Tech, the selection program and much of the analyses reported herein would have been impossible.

## Supplementary information

**Figure S1.**
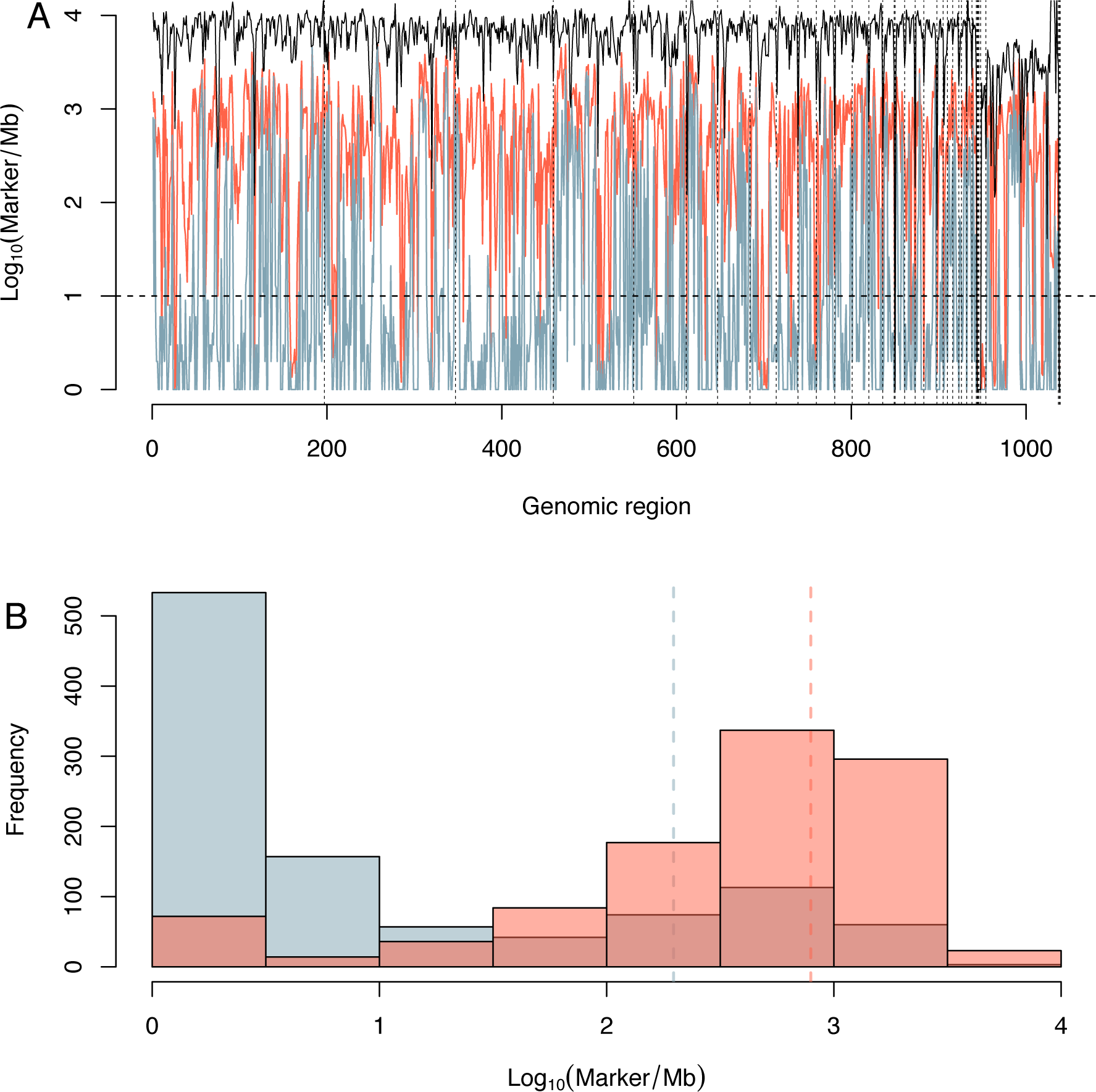
Distribution of the founder line informative SNP markers on chromosome 1. **A)** Line graph illustrating the number of markers in non-overlapping 1Mb bins along chromosome 1 (y-axis; log_10_ transformed). The black/blue/red lines represent the number of segregating markers/markers fixed for alternative alleles in the two founder lines/average number of informative markers within the 73 full-sib F_0_-F_2_ families, all in the Virgina chicken line F_2_ pedigree. **B)** Histograms illustrating the density (x-axis; log_10_ transformed) of informative markers in the Virginia chicken line F_2_ pedigree, where blue/red represent the density markers fixed between the two founder lines/the average marker density of markers fixed for alternative alleles within the 73 F_2_ full-sib families.

**Figure S2.**
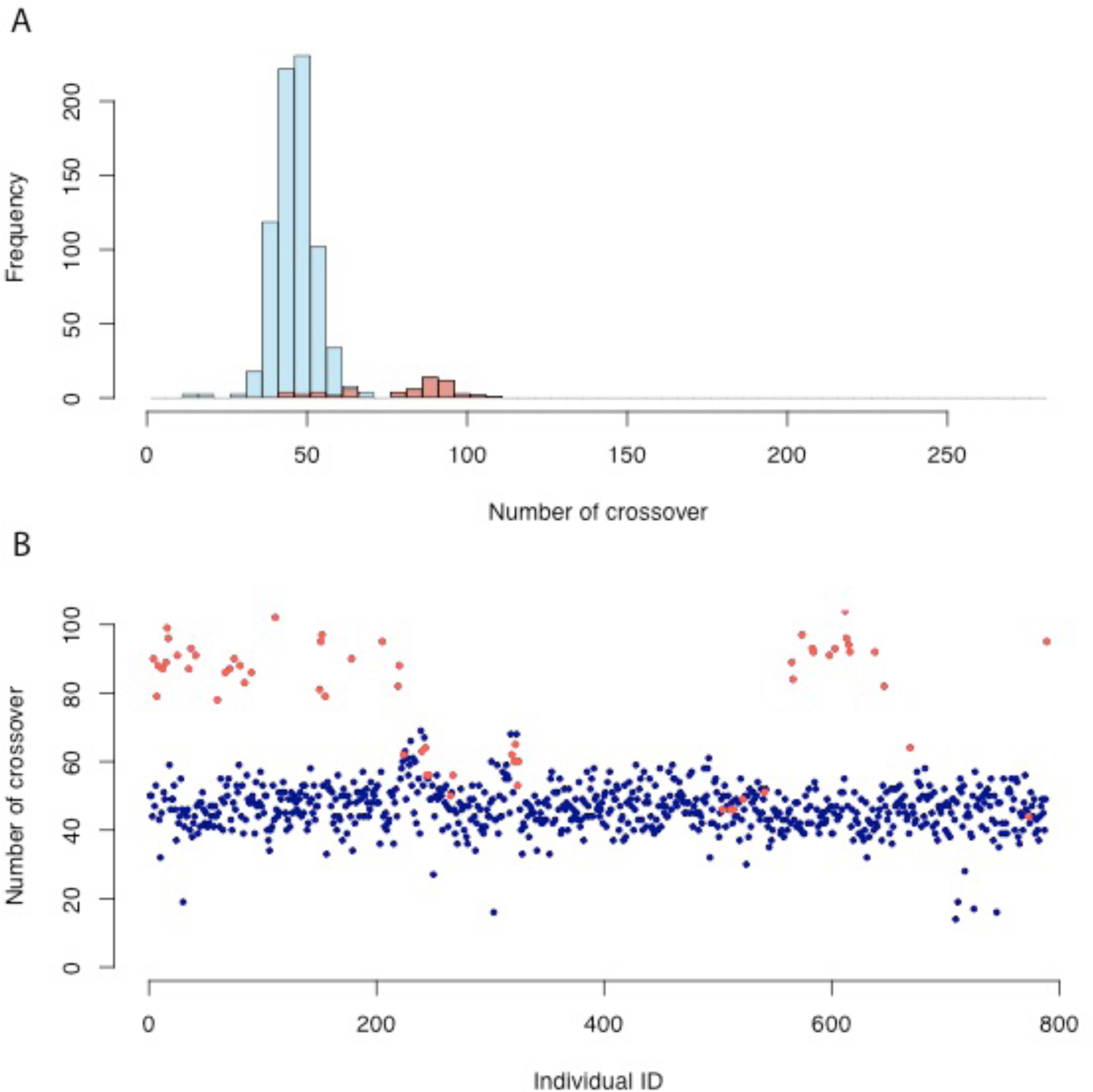
Visualization of the number of crossovers in all F2 individuals. A) Histogram of number of imputed crossover events for the 803 genotyped F_2_individuals; B) Number of imputed crossover events in each individual, sorted by the 73 full-sib families. Individuals with low call rate (call rate <0.9) are colored in red.

**Figure S3.**
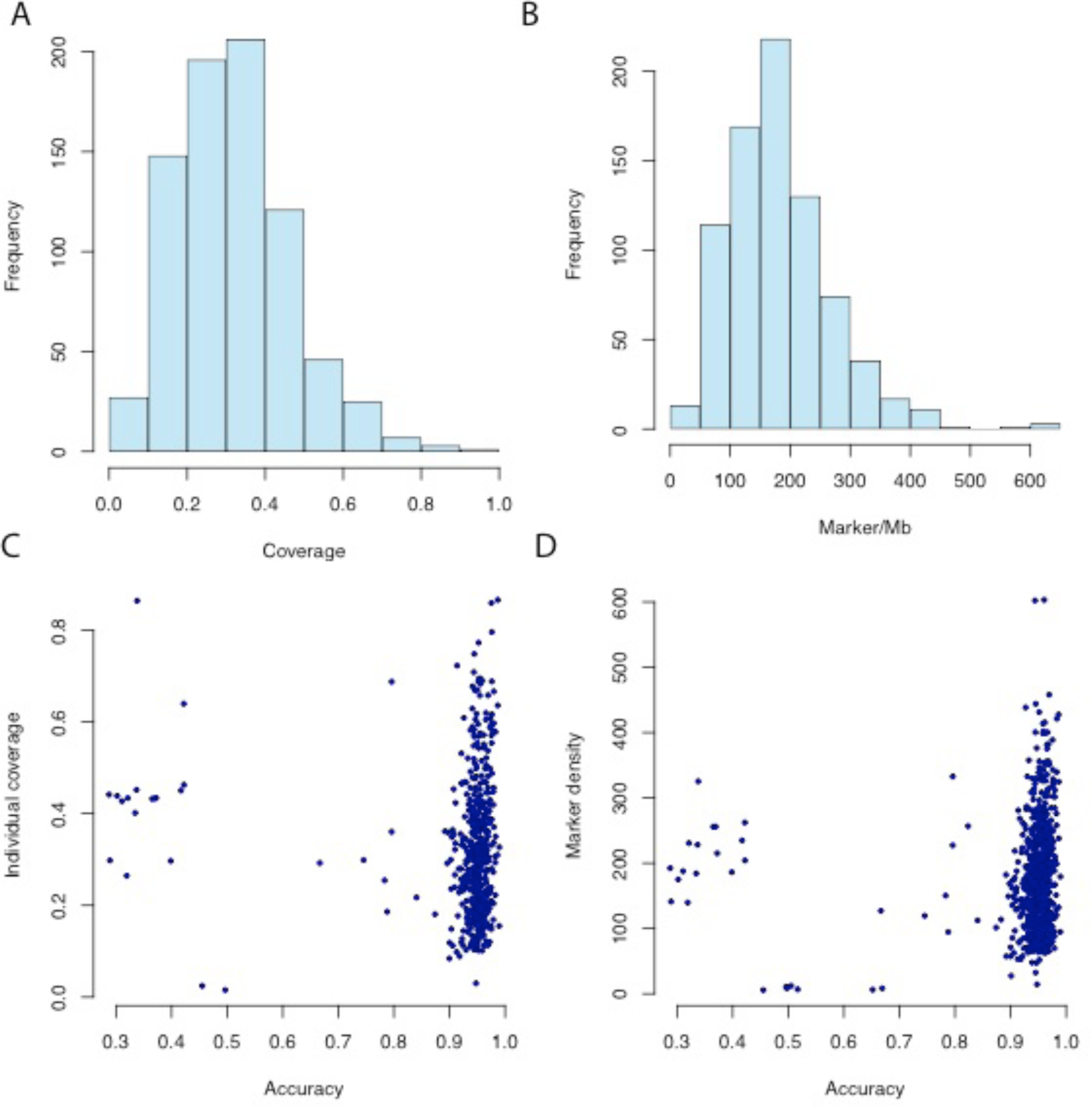
Ilustration of the relationship between sequencing coverage, marker density, and imputation accuracy. **A/B)** Histograms of the sequencing coverage/marker densities for the 803 genotyped F_2_individuals. **C/D)** Scatter plots of individual coverage/marker density vs imputation accuracy measured as the proportion of sites that have same genotype with the averaged genotype probabilities estimated by (Wahlberg et al., 2009) using genotypes of 434 SNP and microsatellite markers with the Haley and Knott algorithm (Haley et al., 1994).

